# Selective endocytic uptake of targeted liposomes occurs within a narrow range of liposome diameter

**DOI:** 10.1101/2023.07.06.548000

**Authors:** Grant Ashby, Kayla E. Keng, Carl C. Hayden, Sadhana Gollapudi, Justin R. Houser, Sabah Jamal, Jeanne C. Stachowiak

## Abstract

Cell surface receptors facilitate signaling and nutrient uptake. These processes are dynamic, requiring receptors to be actively recycled by endocytosis. Due to their differential expression in disease states, receptors are often the target of drug-carrier particles, which are adorned with ligands that bind specifically to receptors. These targeted particles are taken into the cell by multiple routes of internalization, where the best-characterized pathway is clathrin-mediated endocytosis. Most studies of particle uptake have utilized bulk assays, rather than observing individual endocytic events. As a result, the detailed mechanisms of particle uptake remain obscure. To address this gap, we have employed a live-cell imaging approach to study the uptake of individual liposomes as they interact with clathrin-coated structures. By tracking individual internalization events, we find that the size of liposomes, rather than the density of the ligands on their surfaces, primarily determines their probability of uptake. Interestingly, targeting has the greatest impact on endocytosis of liposomes of intermediate diameters, with the smallest and largest liposomes being internalized or excluded, respectively, regardless of whether they are targeted. These findings, which highlight a previously unexplored limitation of targeted delivery, can be used to design more effective drug carriers.

## Introduction

Clathrin-mediated endocytosis is the major uptake pathway of many membrane-bound receptors that initiate signaling^1–3^, interact with the extracellular environment^4^, and regulate nutrient uptake^5, 6^. A nascent clathrin-coated structure is formed when trimers of clathrin, known as triskelia, assemble into an icosahedral lattice on the inner surface of the plasma membrane. Assembly of clathrin causes the membrane surface to bend inward, creating an invagination ^7–9^. Clathrin is recruited to the plasma membrane by a family of adaptor proteins, which also bind to membrane lipids and diverse transmembrane proteins, including most receptors^7, 10, 11^. The endocytic protein network grows as more adaptors and clathrin triskelia are recruited. The resulting network is then able to bend the membrane towards the cytoplasm, creating a clathrin-coated structure^12, 13^. This process continues until a complete clathrin cage is formed around the nascent vesicle, after which the dynamin GTPase cleaves the membrane neck, allowing a vesicle to be internalized into the cell cytoplasm^14, 15^.

Due to their continual uptake from the surfaces of cells, receptors are often the target of drug-carrier particles, such as synthetic liposomes^16, 17^, dendrimers^18, 19^, and inorganic nanoparticles^20, 21^, which can be decorated with ligands that bind to specific receptors. Many distinct receptor species are internalized by clathrin-mediated endocytosis and have therefore been targeted for particle-based delivery. These include receptor tyrosine kinases, G-protein coupled receptors, the transferrin receptor, and the low density-lipoprotein receptor, among many others^2, 22–29^.

While it is well established that many drug-carrier particles are taken into the cell by clathrin-mediated endocytosis, most studies have focused on bulk uptake assays such as flow cytometry^30, 31^ and western blot analysis^32, 33^. These assays, while widely available and relatively straightforward, do not provide insights into the detailed, molecular-scale mechanisms of particle internalization. In contrast, studies focused on the basic science of endocytosis have used microscopy techniques with high spatial and temporal resolution to characterize the dynamic assembly of clathrin-coated structures at the molecular level. In particular, TIRF (total internal reflection fluorescence) microscopy has been used to track the assembly, maturation, and departure of individual clathrin-coated structures at the plasma membrane surface of adherent mammalian cells^8, 34^. This technique only excites fluorophores within about 200 nm of the coverslip surface, facilitating high-resolution imaging of the plasma membrane, with minimal background signal from the cytosol^35^. Using this approach, cell biologists have observed the assembly of individual endocytic structures with high spatio-temporal resolution^8, 34, 36–39^.

Leveraging this approach, we set out to observe the clathrin-mediated internalization of individual targeted liposomes, a class of model drug-carriers^40^. We observed thousands of internalization events, quantifying the impact of targeted liposomes on the probability and dynamics of clathrin-mediated internalization. Importantly, by tracking the uptake of individual liposomes, we were able to distinguish variations in internalization as a function of vesicle size, which is inherently heterogeneous across populations of liposomes and many other types of nano-particle-based carriers^41–43^.

Interestingly, our data demonstrate that penetration of liposomes between and beneath adherent cells constituted a physical barrier, which effectively excluded larger liposomes. This barrier is partially circumvented by targeting, such that larger liposomes penetrated beneath cells when they incorporated targeting ligands. However, targeting failed to significantly increase the fraction of endocytic events that successfully internalized a liposome. This effect was largely explained by the larger size, on average, of the targeted liposomes that penetrated beneath the cell. Specifically, the probability of internalization fell strongly with increasing liposome diameter for both targeted and untargeted liposomes, such that liposomes with diameters of more than 50 nm were rarely internalized by the clathrin pathway. Conversely, very small liposomes, with diameters below 30 nm, were internalized efficiently, whether they were targeted or not. Interestingly, when internalization of liposomes with intermediate diameters of 35-45 nm was evaluated, targeting resulted in a significant increase in uptake. Taken together, these data suggest that targeting liposomes to cell surface receptors can promote penetration of liposomes between cells, a key step toward tissue penetration. However, the selectivity of targeting is optimized within a narrow, intermediate range of liposome diameter. These insights can be used to optimize the efficiency of particle-based therapeutic delivery.

## Results and Discussion

### Targeting enables liposomes to penetrate beneath adherent cells

To study the uptake of individual liposomes by clathrin-coated structures, we needed to target a receptor that is robustly internalized through the clathrin pathway. We chose the transferrin receptor (TfR), which has a strong affinity for clathrin-coated structures, independent of ligand binding^44, 45^. TfR’s cytosolic domain contains a YXXФ motif, which binds adaptor protein 2 (AP2), a major constituent of clathrin-coated structures^22, 46^. We created a chimeric version of TfR in which the extracellular domain was replaced by a monomeric streptavidin domain^47^ and a monomeric eGFP to report the receptor expression level. The monomeric streptavidin domain has a dissociation constant for biotin that is in the nanomolar range, similar to many native ligand-receptor interactions^48, 49^. Our reasons for creating a chimeric receptor, rather than using a native one, were twofold: (i) to be able to precisely monitor the relative expression level of the receptor between cells in live cell imaging experiments of eGFP, and (ii) to be able to use a small molecule, biotin, as the targeting ligand, rather than a macromolecular ligand, which would add significantly to liposome size. In this way, we are able to largely separate the impact of liposome size and the density of the targeting ligand, as illustrated below.

Using this system, we targeted liposomes to cells expressing the chimeric receptor simply by including biotinylated lipids in the membrane composition. The chimeric receptor, TfR-mEGFP-mSA, is shown in Figure 1A. It consists of the cytosolic and transmembrane domains of the transferrin receptor fused to monomeric-streptavidin, followed by an eGFP domain for visualization during live cell imaging. This chimeric receptor was expressed in SUM159 cells that were gene-edited to include a HaloTag in the AP2-σ2 subunit, a generous gift of the T. Kirchhausen laboratory^50^.This tag enables labelling of clathrin structures upon addition of the membrane permeable HaloLigand, JF646.

**Figure 1.**
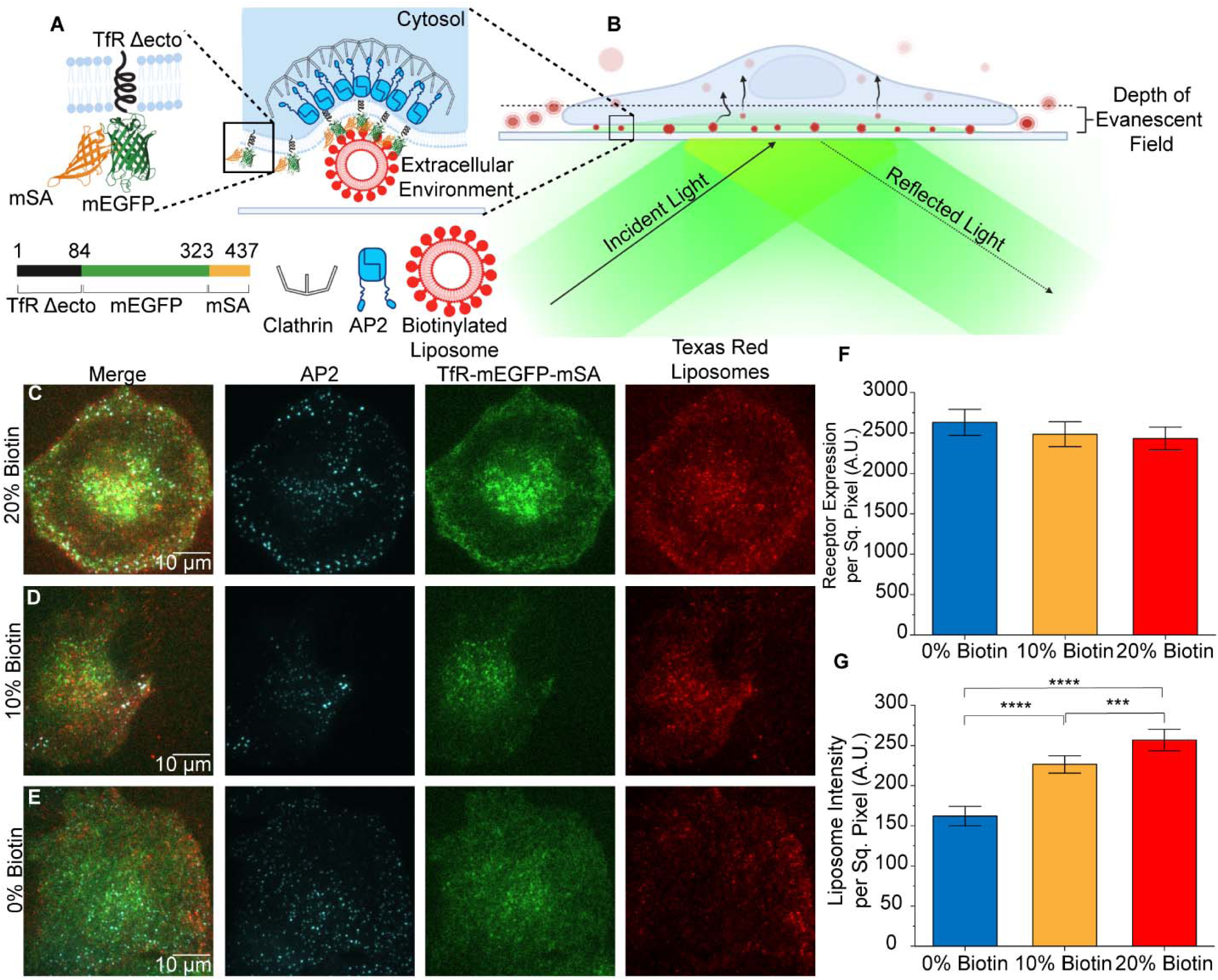
Targeting promotes penetration of liposomes beneath adherent cells. A. The chimeric receptor, TfR-mEGFP-mSA, which was designed to recruit biotinylated liposomes to endocytic sites. **B.** Schematic of TIRF microscopy to examine internalization of liposomes by endocytosis from the adherent surfaces of cells. **C-E.** TIRF microscopy images of AP2 (JF646), chimeric receptor (EGFP), and liposomes (Texas Red DHPE) at the adherent surfaces of SUM159 cells. All channels have been contrasted equally across all three conditions. **F.** Average fluorescence intensity of the chimeric receptor on the plasma membrane. No significant differences were seen between groups of cells exposed to each population of liposomes. (t-test 0% vs 10%; p = 0.514; n = 107, 182) t-test 0% vs 20%; p = 0.353; n = 107; 166; t-test 10% vs 20%; p = 0.800; n = 182, 166). **G.** Total fluorescence intensity per area of liposomes present beneath cells. (t-test 0% vs 10%; p < 1×10^-4^; n = 107, 182) t-test 0% vs 20%; p < 1×10^-4^; n = 107; 166; t-test 10% vs 20%; p = 3.55×10^-4^; n = 182, 166). For **F** and **G**, three independent trials were acquired for each condition with a minimum of 15 cells imaged per trial. Error bars represent the standard error of the mean, where N is the number of cellular crops analyzed across all trials. Significance between conditions was identified using a two-tailed student’s t-test and a one-factor ANOVA with α = 0.05.

Liposomes contained 0 – 20 mol% of biotinylated lipids (PE-CAP-biotin), along with 10 mol% Texas Red-DHPE for visualization, and 2 mol% of pegylated lipids to reduce non-specific binding (DSPE-PEG2K). The remaining portion of each liposome consisted of DOPC. The distribution of liposome diameters, which averaged 70 to 80 nm, was quantified using dynamic light scattering, which indicated that incorporation of biotinylated lipids did not cause a systematic shift in liposome size (Supplemental Figure 1).

To visualize interactions between liposomes and clathrin-coated structures, we employed total internal fluorescence (TIRF) microscopy. TIRF illumination restricts the excitation of fluorophores to a region within 100-200 nm from the top surface of the coverslip. This approach is ideal for isolating the plasma membrane from background fluorescence in the cellular cytosol (Figure 1B). Using TIRF microscopy to image cells that expressed the chimeric receptor, we observed interactions between clathrin-coated structures and liposomes. Cells were exposed to liposomes containing either 0, 10, or 20 mol% biotinylated lipids, 15 minutes prior to imaging (1 μM total lipid). Interestingly, as the percentage of biotinylated lipids in the liposomes increased, the total intensity of the liposomes that penetrated beneath the cell also increased (Figure 1C-E), suggesting that the targeting ligand enhanced the penetration of liposomes beneath cells and into the TIRF field.

To evaluate this result more quantitatively, we used open-source analysis software by Aguet et. al. to detect diffraction-limited puncta in the Texas Red DHPE channel, which represented the liposomes^51^. Specifically, we summed the intensity of these puncta per area beneath the cell, comparing groups of cells that expressed similar levels of the chimeric receptor (Figure 1F). These data confirmed that the intensity associated with liposomes beneath cells was approximately 46% higher for cells exposed to liposomes that contained 20 mol% of biotinylated lipids than for cells exposed to liposomes that did not contain biotinylated lipids (Figure 1G).

## Liposomes associate with clathrin-coated structures that have longer lifetimes

We next sought to determine how internalization of targeting vesicles impacts the dynamics of clathrin-coated structures. Importantly, when liposomes are internalized by clathrin-coated structures, both the liposome and the clathrin-coated structure must exit the narrow region of TIRF illumination, such that they should disappear from TIRF images. Because liposomes are otherwise trapped beneath cells, total loss of liposomal intensity, provided it occurs well away from the edge of the cell, can be confidently interpreted as internalization of the liposome. Further, if this loss of intensity is strongly correlated with the presence and subsequent loss of a punctum in the AP2 fluorescence channel (JF646), we can conclude that internalization of the liposome likely occurred through the clathrin pathway. Leveraging these advantages, we investigated the impact of liposomes on the dynamics of the clathrin-coated structures that associated with them.

Clathrin coated structures assemble and mature at the plasma membrane surface before departing into the cytosol, as discussed above. The assembly process is highly heterogeneous, lasting from ten seconds to several minutes. Not all assemblies of endocytic proteins lead to productive vesicles. As many as half of all endocytic assemblies stochastically abort without creating a vesicle^15, 52^. Endocytic assemblies that remain at the plasma membrane for less than 20 seconds are likely to have aborted, based on imaging studies in which markers of vesicle scission were tracked^15^. While many assemblies with lifetimes longer than 20 seconds lead to productive endocytosis, clathrin-coated structures that remain at the plasma membrane for longer than several minutes often represent stalled endocytic events that do not lead to internalization and may ultimately be removed from the cell surface by autophagy^53^. Here we sought to determine how the native dynamics of clathrin-coated structures are impacted by internalization of targeted liposomes.

To observe the impact of targeted vesicles on the dynamics of clathrin-coated structures, we added liposomes to SUM159 cells that expressed the chimeric receptor. A representative TIRF image of Texas Red labeled liposomes (red) interacting with clathrin-coated structures, as marked by AP2 (JF646 shown in cyan) is shown in Figure 2A. Colocalization of puncta in the liposome and AP2 channels indicates interaction between liposomes and clathrin-coated structures. We recorded the fluorescence intensity of colocalized, diffraction-limited puncta over time, as shown in Figure 2B. The simultaneous drop in the fluorescence intensity of both the clathrin coated structure and the liposome indicated successful internalization of a liposome by clathrin-mediated endocytosis (Figure 2C). Using this approach, we utilized the same openly available analysis package, CMEanalysis, to track thousands of colocalization events across tens of cells^51^.

**Figure 2.**
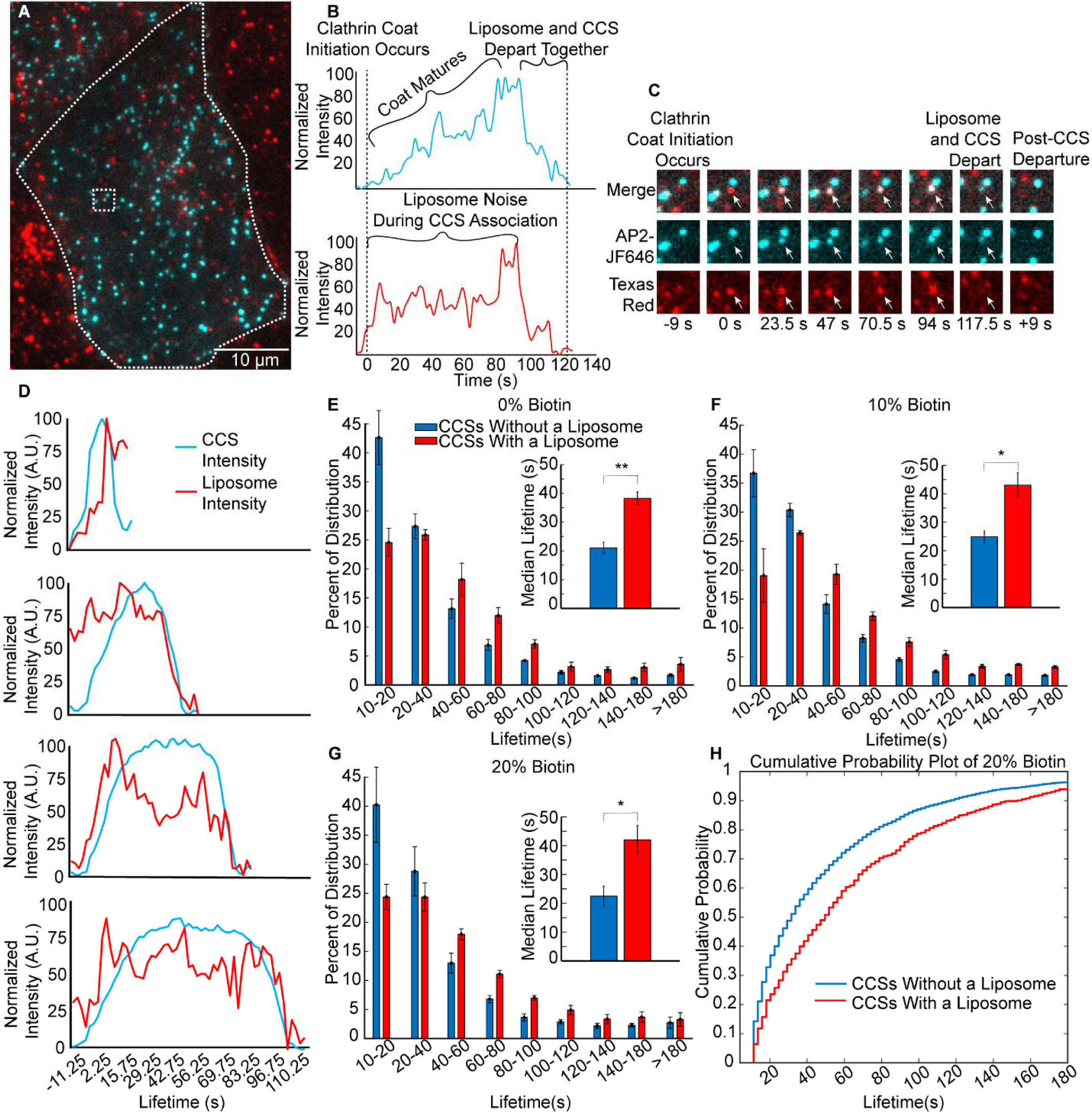
Liposomes associate with clathrin-coated structures that have longer lifetimes. A. A TIRF microscopy image at the plasma membrane of a SUM159 cell, gene edited to express a HaloTag on the σ-subunit of AP2, incubated with 1μM of liposomes (total lipid), which contained 10 mol% of biotinylated lipids. **B.** Fluorescence intensity as a function of time for an individual liposome, which colocalized with an individual clathrin-coated structure. The liposome and clathrin-coated structure decay in intensity over the same period of time, suggesting simultaneous departure from the TIRF field, as expected for internalization of a liposome by endocytosis. **C.** Montage of images from **B**, where the white arrow indicates the tracked structure. **D.** Average fluorescence intensity over time for endocytic structures with lifetimes within the following ranges: 10 to 2 s, 40 to 60 s, 60 to 80 s, and 80 to 100 s. Intensity shown for the liposome (Texas Red DHPE) and AP2 (JF646) channels. Cohorts were composed of 1647, 739, 563, and 368 events, for the 10-20s, 40-60s, 60-80s, and 80-100s cohorts, respectively. **E-G.** Distribution of clathrin-coated structure lifetimes for cells exposed to liposomes containing: 0 mol% (**E**), 10 mol% (**F**), and 20 mol% (**G**) biotinylated lipids. The insets of parts **E-G** compare median lifetimes of the clathrin structures that were associated with a liposome to those that did not. Error bars represent the standard error of the mean of N = 3 independent trials. Total number of clathrin-mediated endocytic events per graph was 8,804, 22,869, and 10,930, respectively. **H.** Cumulative probability of endocytic structure departure as a function of time for the data shown in **G**.

We then filtered out clathrin-coated structures that interacted for a significant period of their lifetime with a liposome (see methods). We grouped the resulting clathrin-coated structures into cohorts based upon their lifetime at the plasma membrane surface, from 10 to 180 seconds. The average intensity over time in the liposome and AP2 channels for several of these cohorts are shown in Figure 2D. As the clathrin-coated structure grows and matures, its intensity gradually increases, reaching a maximum value before disappearing from the TIRF field, as indicated by the rise and subsequent fall in the intensity of the AP2 signal (JF464), shown in cyan in Figure 2D. The liposomal signal does not necessarily match the initial rise in the AP2 signal, because liposomes are typically present in the optical plane prior to internalization. However, the simultaneous decline in the intensity of the AP2 and liposome channels indicates internalization of a liposome, Figure 2B, C. In the 10 – 20 second cohort, which contains the shortest-lived clathrin-coated structures, the liposome signal did not drop with the AP2 signal, indicating that most of the structures within this cohort failed to internalize a liposome, likely because they were abortive^15, 52^. In contrast, a simultaneous decay in AP2 and liposome intensity was observed for cohorts that contained longer-lived structures, for example, 40–60, 60–80, and 80-100 seconds, Figure 2D. These data suggest that liposomes are successfully internalized by clathrin-coated structures with a diverse range of lifetimes.

To examine the impact of liposomes on the dynamics of clathrin-coated structures, we plotted the distribution of lifetimes for clathrin-coated structures within cells exposed to liposomes with 0, 10, or 20 mol% biotinylated lipids, Figure 2E-G. In each graph, the fraction of clathrin-coated structures within each of the temporal cohorts is plotted as a series of bars. The data were divided into two subsets: structures that did not colocalize with a liposome (blue bars), and structures that did colocalize with a liposome (red bars). The summation of these groups equates to the corresponding curve for the full population of endocytic structures, shown in Supplementary Figure 2. Our data show that for each group of liposomes, whether targeted or untargeted (Figure 2E), the clathrin-coated structures that associate with a liposome tend to be longer-lived than those structures that do not associate with liposomes. This trend could occur for one of two reasons: (i) the presence of liposomes stabilizes endocytic structures, preventing them from aborting, or (ii) the longer a clathrin-coated structure resides at the membrane, the higher the probability that a liposome will interact with it. To distinguish between these possible explanations, we examined the impact of liposomes on the overall distribution of lifetimes for all endocytic structures (Supplementary Figure 2). This distribution, which contains structures that associated with liposomes, as well as those that did not, was not substantially shifted from the corresponding distribution for endocytic structures within cells that were never exposed to liposomes. Based on these data, it appears unlikely that liposomes stabilize endocytic sites. Instead, it appears that liposomes interact more with endocytic structures that reside for longer times at the plasma membrane. This trend is summarized in Figure 2H, which compares the cumulative probability that an endocytic structure will depart from the plasma membrane as a function of time, for the populations of structures that do and do not associate with liposomes, respectively. From these data it is evident that liposomes associated with a population of endocytic structures have longer than average lifetimes at the plasma membrane, likely because the longer lifetime increases the probability that a liposome will encounter an endocytic structure before it disappears from the cell surface. Notably, our analysis so far has concentrated on the impact that association with a liposome has on the dynamics of endocytic structures. Of those endocytic structures that associate with a liposome, only a fraction will successfully internalize it. In the next section we examine the impact of targeting on the fraction of associations that progress to internalization.

### Targeting does not significantly impact the overall probability that a liposome will be internalized by a clathrin-coated structure

Having established that liposomes, whether targeted or untargeted, have a minimal effect on endocytic dynamics, we next investigated the impact of targeting on the probability of liposome internalization. Figure 3A shows the fraction of liposomes that associated with clathrin-coated structures, as a function of the biotinylated lipid content of the liposomes. Surprisingly, these data suggest that targeting, via inclusion of biotinylated lipids, did not increase the fraction of liposomes that associated with endocytic structures. We next asked if targeting impacted the fraction of associated liposomes that were successfully internalized by endocytic structures. For this purpose, it was necessary to identify bona fide internalization events within our data set. Such events were characterized by the simultaneous disappearance of the fluorescence signal in the liposome (Texas Red) and AP2 (JF646) channels, as described under materials and methods. Figure 3B plots the number of liposome internalization events per membrane area for cells exposed to liposomes of increasing biotinylated lipid content. Interestingly, these data indicate that the probability of internalization by an endocytic structure is largely independent of biotin content, similar to the probability of association to endocytic structures (Figure 3A).

**Figure 3.**
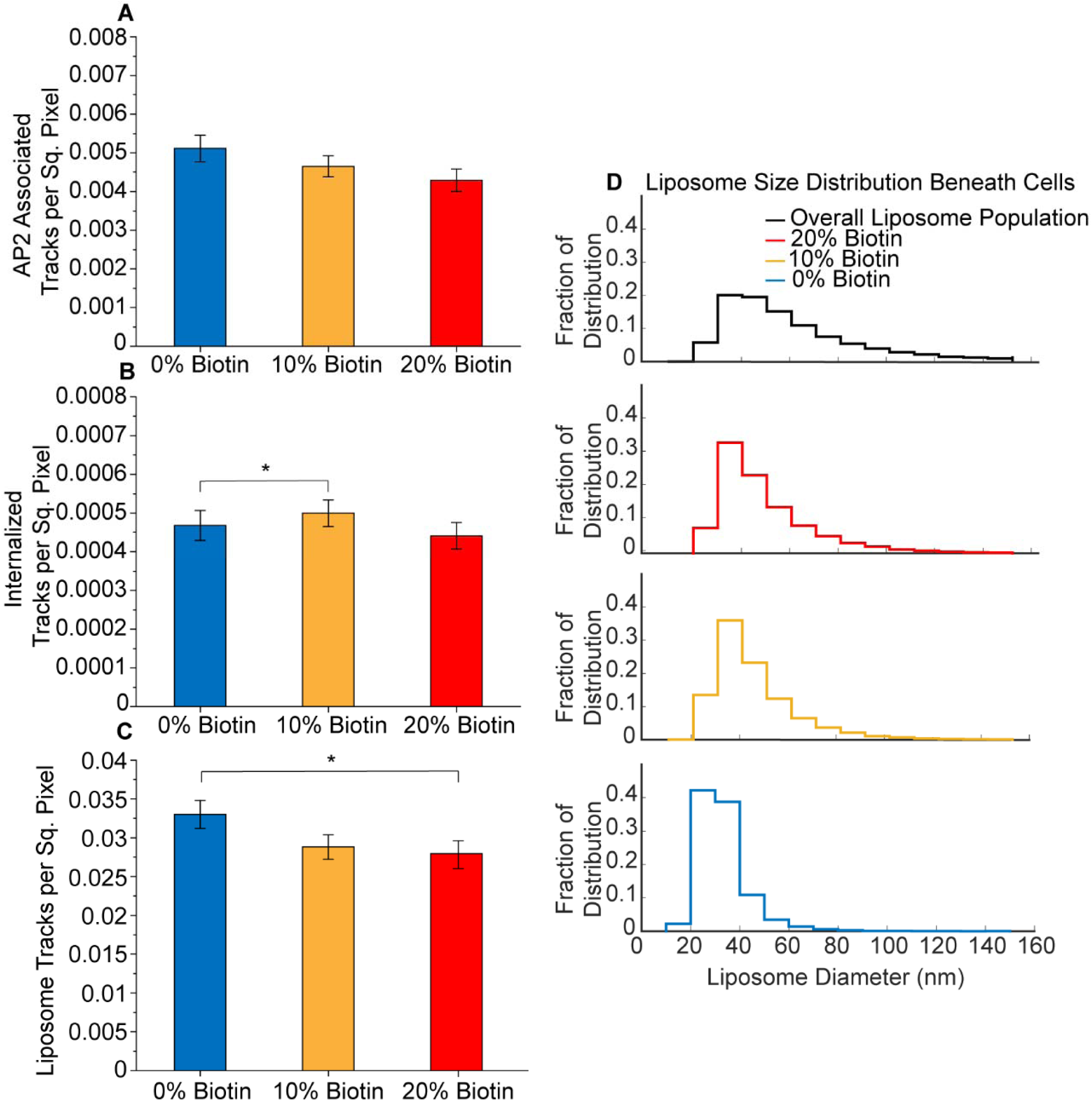
Targeting does not significantly impact the overall probability that a liposome will be internalized by a clathrin-coated structure. A. Bar graph of the number per area of liposomes that associated with a clathrin-coated structure for liposomes containing 0, 10, and 20 mol% of biotinylated lipids (t-test 0% to 10%; p = 0.070; n = 66, 84; t-test 0% to 20%; p = 0.523; n = 66, 68; t-test 10% to 20%; p = 0.256; n = 84, 68). **B.** Bar graph of the number per area of liposomes that were in internalized by a clathrin-coated structure for liposomes containing 0, 10, and 20 mol% of biotinylated lipids (t-test 0% to 10%; p = 0.082; n = 66, 84; t-test 0% to 20%; p = 0.048; n = 66, 68; t-test 10% to 20%; p = 0.722; n = 84, 68). **C)** Bar graph of the number per area of total liposomes beneath cells for liposomes containing 0, 10, and 20 mol% biotinylated lipids (t-test 0% to 10%; p = 0.015; n = 66, 84; t-test 0% to 20%; p = 0.417; n = 66, 68; t-test 10% to 20%; p = 0.124; n = 84, 68). For **A-C**, three independent trials were acquired for each condition with a minimum of 15 cells imaged per trial. Cells across trials were combined for a total of 66, 84, and 68 cellular crops exposed to liposomes containing 0, 10, and 20 mol% biotinylated lipids, respectively. Error bars represent the standard error of the mean. No significance between conditions was identified using a two-tailed student’s t-test with α = 0.05. **D.** Distribution of liposome diameters. The black curve is the overall size distribution of liposomes prior to their exposure to cells (47,502 liposomes). The red, gold, and blue curves are the size distributions for liposomes that penetrate beneath cells, for liposomes containing 20, 10, or 0 mol% biotinylated lipids. The red, gold, and blue distributions contain 56,130, 61,002, and 31,150 liposomes respectively.

How can we reconcile the observation that targeting increases penetration of liposomes beneath adherent cells (Figure 1G), with the seeming inability of targeting to drive an increase in internalization of liposomes by endocytosis (Figure 3A,B)? Toward answering this question, we probed deeper into the results in Figure 1G. In particular, there are two possible explanations for the increase in liposome intensity beneath the cell with increasing concentration of the targeting ligand: (i) targeting increases the number of liposomes that penetrate beneath cells, or (ii) targeting increases the size of the liposomes that penetrate beneath cells. Notably, these explanations are not mutually exclusive. To determine their relative role in explaining the trends in Figure 1G, we began by counting the number of liposomes per area beneath the cell as a function of the concentration of the targeting lipid (Figure 3C). Here we find no increase in liposome number with increasing biotin content. Therefore, we next examined the distribution of diameters for liposomes present beneath cells.

For this purpose, we used an intensity-based analysis to determine a conversion factor between the diameter of a liposome and the brightness of the fluorescent puncta it creates in TIRF images. Using this approach, which we have previously reported^54^ the distribution of diameters for liposomes tethered to a coverslip could be approximated, as shown in Figure 3D (black curve). These data represent the initial distribution of liposome diameters prior to their exposure to cells. When this approach was applied to liposomes present beneath cells, the distribution of diameters shifted towards smaller values, suggesting that larger liposomes are less likely to penetrate beneath adherent cells (Figure 3D, blue curve). The addition of biotinylated lipids (gold and red curves) partially overcame this limitation, allowing a higher fraction of larger liposomes to penetrate. The larger size of liposomes present beneath cells explains the increase in the fluorescent intensity of the liposomes in Figure 1G. It may also explain the failure of targeting to substantially increase the probability that liposomes associate with and become internalized by endocytic structures, Figure 3A, B. Specifically, previous work has suggested that larger particles are internalized less efficiently by the clathrin pathway^55–57^.

### Targeting increases the probability of endocytic uptake for liposomes of intermediate diameter

Having established that inclusion of biotinylated lipids enables larger liposomes to penetrate beneath cells, we next examined the impact of a liposome’s size on its probability of internalization, comparing targeted and non-targeted liposomes. Figure 4A-D compares the distribution of liposome diameters for four cases: (A) liposomes prior to interaction with cells (repeated from Fig. 3D), (B) liposomes that penetrate beneath the cell (repeated from Fig. 3D), (C) liposomes that penetrate beneath the cell and associate with an endocytic structure, and (D) liposomes that penetrate beneath the cell, associate with an endocytic structure, and become internalized. In each case, data are compared for liposomes that lacked biotinylated lipids (blue curves) and those that contained 20 mol% of biotinylated lipids (red curves). As described above, exclusion of non-targeted liposomes from the space beneath the cell shifts the distribution of diameters toward smaller values (compare Figure 4A to blue curve in B). In contrast, targeted liposomes beneath the cell have diameters that more closely mimic the initial size distribution of the liposomes, prior to their exposure to cells (compare Figure 4A to red curve in B). Interestingly, when we examine liposomes that associate with endocytic structures, the distribution of liposome diameters shifts toward smaller values for both targeted and non-targeted liposomes, such that there is little difference between the two distributions, Figure 4C. This result suggests that smaller liposomes are more likely to find developing endocytic structures, perhaps owing to increased mobility within the very limited space between the coverslip and the adhered cell. Similarly, if we examine liposomes that are internalized by endocytic structures, a subset of those that are associated, we again find that smaller liposomes are more likely to be internalized and that the distribution of diameters for internalized liposomes differs little between targeted and non-targeted liposomes, Figure 4D.

**Figure 4.**
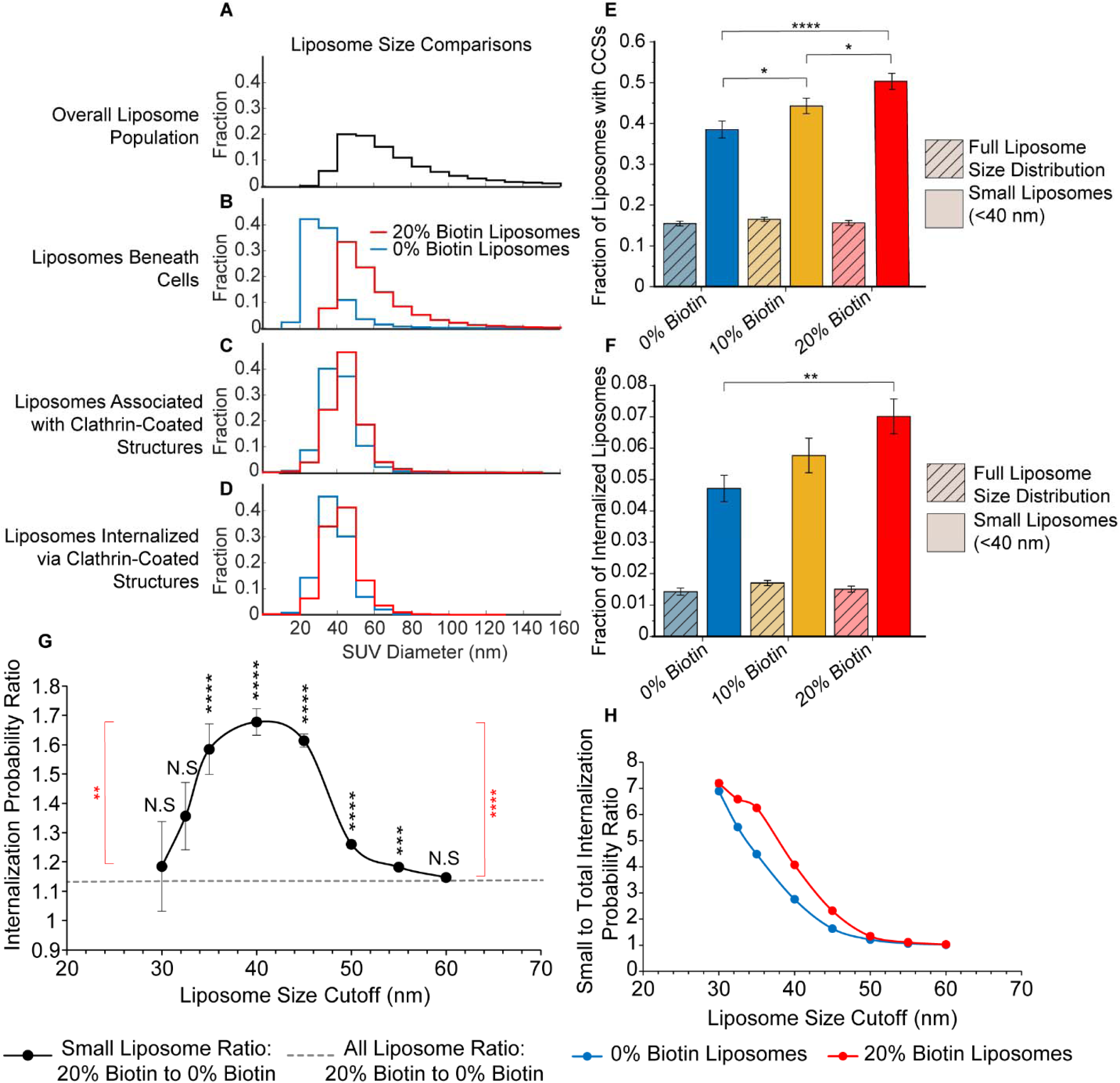
Targeting increases the probability of endocytic uptake for liposomes of intermediate diameter. A. Distribution of liposome diameters for liposomes containing 0% or 20% biotinylated lipids. The top plot (black curve) is the overall distribution of liposome diameters (n = 47,502), repeated from Figure 3. The second plot is the distribution of diameters for liposomes that penetrated beneath cells (repeated from Figure 3). The third plot is the distribution of diameter for liposomes that penetrated beneath cells and associated with a clathrin-coated structure (12,282 (red) and 16,821 (blue) liposomes). The fourth plot is the distribution of diameters for liposomes that penetrated beneath cells, associated with a clathrin-coated structure, and became internalized (1,269 (red) and 1,577 (blue) liposomes). **B.** Bar graph representing the probability that a liposomes will associated with a clathrin-coated structure for the full distribution of liposome diameters (hashed bars) and the population of liposomes with diameters below 40 nm (solid bars), for liposomes containing 0 (blue), 10 (gold), or 20 (red) mol% biotinylated lipids (t-test 0% to 10%; p = 0.042; n = 66, 84; t-test 0% to 20%; p < 1×10^-4^; n = 66, 68; t-test 10% to 20%; p = 0.028). **C.** Bar graph representing the probability that a liposomes will be internalized by a clathrin-coated structure for the full distribution of liposome diameters (hashed bars) and the population of liposomes with diameters below 40 nm (solid bars), for liposomes containing 0 (blue), 10 (gold), or 20 (red) mol% biotinylated lipids (t-test 0% to 10%; p = 0.133; n = 66, 84; t-test 0% to 20%; p = 0.001; n = 66, 68; t-test 10% to 20%; p = 0.115). For **B and C**, N = 3 independent trials were run for each condition, with at least 15 cells imaged per trial. The total number of cellular crops from all trials in **B-C** was 66, 84, and 68 for liposomes containing 0, 10, or 20 mol% biotinylated lipids, respectively. Error bars represent the standard error of the mean for each population. Significance between conditions was identified using a two-tailed student’s t-test with α = 0.05. **D.** The ratio of the probability of internalization for liposomes containing 20 versus 0 mol % biotinylated lipids, plotted as a function of the liposome diameter cutoff. The average ratio for the entire population of liposome diameters is shown in orange. The significance of the differences between the data for each cutoff (black) and the population average (orange) were determined using a z-test (two-sample for means; 30 nm p = 0.748; 32.5 nm p = 0.055; 35 nm p = < 1×10^-5^; 40 nm p < 1×10^-5^; 45 nm p < 1×10^-5^; 50 nm p = < 1×10^-5^; 55 nm p = 0.000393; 60 nm p = 0.371; n = 82, n = 82). **E.** The ratio of the probability of liposome internalization for liposomes with diameters below a specific cutoff (horizontal axis) relative to the overall probability of internalization for the full population of liposomes (all diameters). The significance of differences between the data for liposomes containing 0 (blue) and 20 mol% (red) biotinylated lipids was determined using a z-test (two-sample for means; 30 nm p =0.0506; 35 nm p < 1×10^-5^; 40 nm p < 1×10^-5^; 45 nm p < 1×10^-5^; 50 nm p < 1×10^-5^; 55 nm 0.000868; 60 nm p = 0.422; n = 82, n = 82). The total number of cells in **D and E** were 82 and 82, respectively. Error bars represent the standard deviation.

These results provide a possible explanation for our finding that the overall probability of liposome internalization is not strongly impacted by targeting (Figure 3A,B). Specifically, while targeting enables larger liposomes to penetrate beneath the cell (Figure 3D), this effect appears to be largely neutralized by the much less efficient internalization of larger liposomes (Figure 4C,D). If so, we would expect liposomes with small diameters to experience the greatest increase in internalization upon targeting.

To test this idea, we compared the efficiency of internalization between targeted and non-targeted liposomes with diameters below 40 nm. This threshold was chosen because it is approximately at the median of diameter distribution for liposomes that penetrated beneath cells, for both the targeted and non-targeted populations (Figure 4B). Figure 4E shows that the frequency with which these small liposomes associated with clathrin-coated structures was higher compared to the overall population, a trend which increased with targeting. Similarly, the frequency of internalization was also greater for liposomes with diameters below 40 nm, Figure 4F.

To further explore the impact of liposome diameter on targeting, we compared the probability of internalization for liposomes containing 20% biotin to the corresponding probability of internalization for untargeted liposomes, using a diameter cutoff that varied from 30 to 60 nm. A ratio of 1 between these probabilities would indicate that there is no difference in internalization due to targeting. Considering the entire population of liposome diameter, without using a cutoff, the ratio was 1.1, indicating that liposomes that contained 20% of biotinylated lipids were only about 10% more likely to be internalized than untargeted liposomes, as shown by the horizontal line in Figure 4G. Examining liposomes with diameters below 32.5 nm, the internalization probability ratio did not differ significantly from 1.1, indicating a lack of selective targeting. However, the internalization probability ratio was significantly higher when the diameter cutoff was between 35 and 55 nm. A threshold of 40 nm resulted in the highest ratio of approximately 1.6, indicating that liposomes containing 20 mol% of the biotinylated lipid were about 60% more likely to be internalized than untargeted liposomes. For cutoffs above 55 nm, the internalization probability ratio was no longer significantly greater than the overall population average, indicating that targeting failed to create selectivity, Figure 4G.

These results are further elucidated in Figure 4H, which plots the relative probability of liposomal internalization below the diameter cutoff on the horizontal axis. Here it is evident that the probability of internalization declines monotonically with increasing liposome diameter, for both targeted (20 mol% biotinylated lipids) and untargeted liposomes, with the smallest liposomes having an uptake probability about 7-fold higher than the average liposome within the population. The smallest liposomes appear to be easily internalized, regardless of targeting, likely because they are highly mobile and too small to sterically interfere with endocytosis. A gap between the curves emerges for liposomes with diameters between 32.5 nm and 50 nm. Within this range, the uptake probability is higher for targeted liposomes. This gap closes for liposomes with diameters greater than 50 nm, which are unlikely to be internalized, regardless of targeting. Poor internalization of these larger liposomes is likely the result of immobility and steric inhibition of endocytosis, as reported previously^42, 55, 58, 59^.

## Discussion

Here we used an *in vitro* targeting system based on a chimeric transmembrane receptor to investigate the impact of targeting on the uptake of liposomes by the clathrin-mediated endocytic pathway. We employed TIRF microscopy to observe interactions between individual liposomes and growing clathrin-coated structures at the plasma membrane surface of adherent mammalian epithelial cells. While established techniques such as flow cytometry and western blot can measure overall cellular uptake of liposomes and other nanoparticles^30–33, 60–64^, these approaches do not permit the observation of individual uptake events, such that it is not possible to study the impact of intrinsic heterogeneity across a population of particles. In contrast, by using TIRF microscopy to achieve real-time monitoring of endocytosis in live cells, we were able to observe thousands of individual internalization events. This approach allowed us to isolate the differential impact of liposome size and targeting on the probability of liposome internalization, factors which have been difficult to deconvolute in previous work.

To our surprise, we found that vesicle size had a much greater impact on the efficiency of liposomal uptake by the clathrin pathway, compared to targeting, Figure 5. Specifically, while targeting substantially increased the size of liposomes that penetrated beneath adherent cells, the inclusion of targeting ligands in vesicles had only a slight impact on the efficiency with which these liposomes were ultimately internalized. In contrast, when vesicles of different sizes were compared within the heterogenous liposome population, the efficiency of internalization was approximately 700% higher for the smallest vesicles (30 nm diameter) compared to the largest vesicles (60 nm diameter), Figure 4H. Within this range, a positive impact of targeting on internalization efficiency was observed only for vesicles within a narrow range of diameters from 35-50 nm, where the maximum magnitude of the increase was 60-70%, Figure 4G. The approximately 10-fold greater impact of liposomes size relative to targeting appears to arise from the greatly reduced ability of larger liposomes to colocalize with transient endocytic events (Figure 4A-D), likely owing to the crowded environment beneath adherent cells. The difficulty that larger liposomes experience in penetrating this space, which is populated by focal adhesions and extracellular matrix components, is in line with established understanding of reduced tissue penetration by larger particles^65–69^.

**Figure 5.**
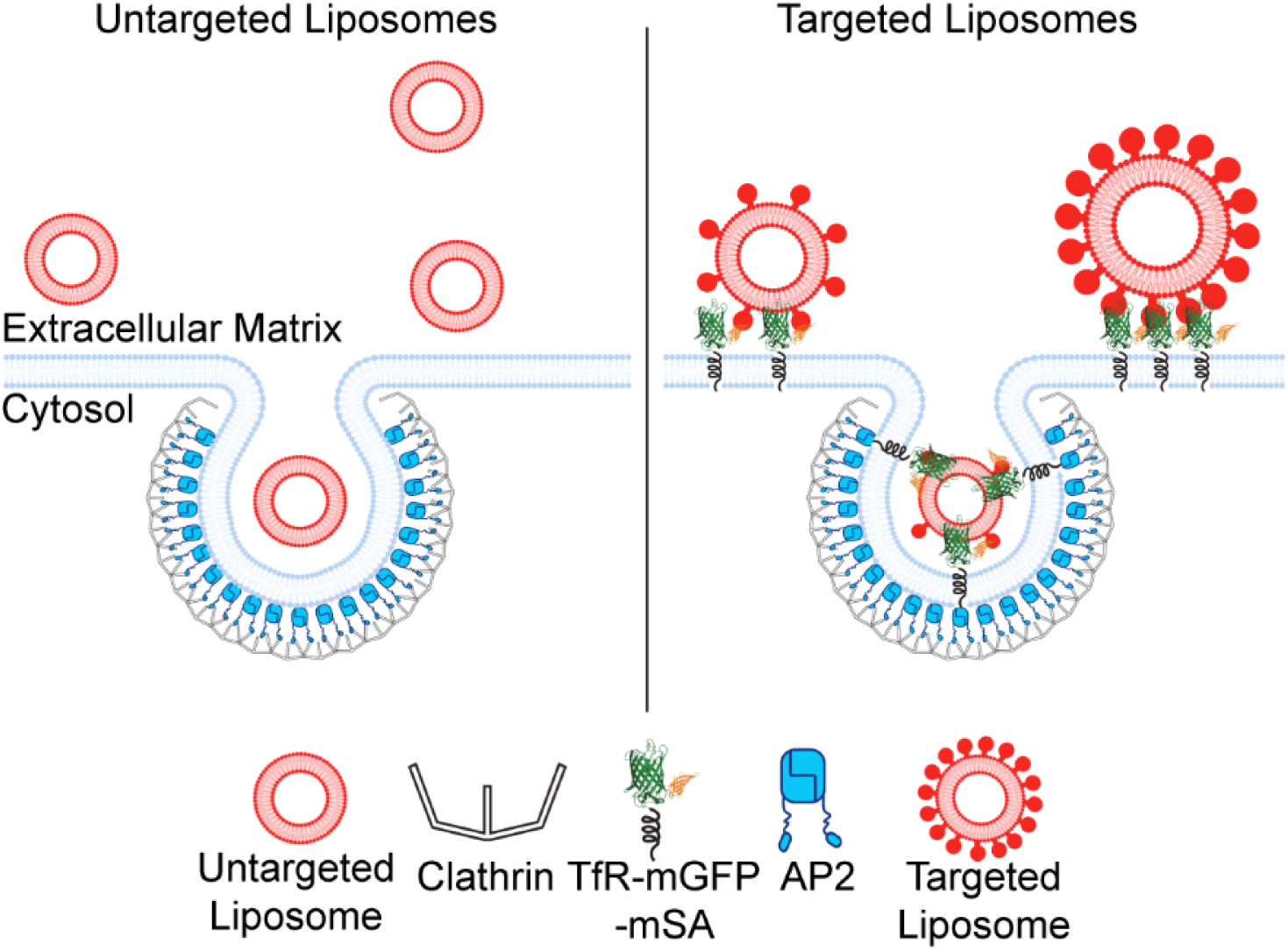
Schematic showing the ability of larger liposomes to penetrate beneath the basolateral cellular membrane due to targeting. In contrast, small liposomes have a high probability of penetration and internalization, regardless of targeting.

Some of the most popular targets for selective drug delivery are receptors that are primarily internalized by the clathrin pathway. These include many nutrient receptors, receptor tyrosine kinases, and G-protein coupled receptors^2, 22–29^. While multiple internalization pathways can play a role in uptake of targeted particles^55, 67, 70, 71^, the clathrin pathway is likely to play an important role in internalization of particles that target these receptors. In this context, our results, which demonstrate that targeting is most selective within a narrow range of particle diameter, suggest a previously unknown “design rule” for targeted particles.

Paradoxically, while inclusion of targeting ligands promoted penetration or larger liposomes beneath cells, the inability of these larger particles to move freely beneath cells prohibited them from being efficiently internalized through the clathrin pathway, Figure 5. While our experiments occurred in a highly simplified in vitro context, they suggest that a similar paradox may occur in vitro, where targeting may improve tissue penetration without resulting in a significant increase in delivery to target cells. This reasoning, along with the many other complexities of delivery in vivo, may help to explain why the success of targeting *in vitro* is often diminished in vivo ^20, 67, 72–74^. Going forward, we anticipate that TIRF-based tracking of individual internalization events can be applied to diverse particle-based delivery systems to gain mechanistic understanding of interactions between engineered particles and the cell’s endocytic machinery.

## Materials and Methods

### Chemical reagents

HEPES, NaCl, Neutravidin, and PLL-PEG (poly-L-Lysine) were purchased from ThermoFisher Scientific. PEG2K-DSPE (1,2-Distearoyl-sn-glycerol-3-phosphorylethanolamine-N [methoxy (polyethylene glycol)-2000]), PE-CAP-Biotin (1,2-dioleoyl-sn-glycerol-3-phosphoethanolamine-[N-(cap biotinyl)]), and DOPC (1,2-Dioleoyl-sn-glycero-3-phosphocholine) were purchased from Avanti Polar Lipids, Inc. Texas Red-DHPE (1,2-dihexadecanoly-snglvero-3-phosphoethanolamine-[N-(Texas Red sulfonyl)]) was purchased from AAT Bioquest. PEG-biotin (Biotin-PEG SVA, MW 5000), and amine-reactive PEG (mPEG-Succinimidyl Valerate, MW 5000) were purchased from Laysan Bio, Inc.

### Plasmids

The plasmid encoding the chimeric receptor (TfRΔecto-mEGFP-mSA) was constructed by inserting mSA and then mEGFP into a pEGFPN1 mammalian expression vector containing TfRΔecto-RFP by Gibson Assembly cloning. First, the mSA gene was inserted downstream of RFP to create TfRΔecto-RFP-mSA. A plasmid encoding pDisplay-mSA (Addgene #39863) was generously provided by Dr. Sheldon Park (University of Buffalo). The mSA fragment was isolated using the forward primer 5’-GCCGCCACTCCACCGGCGCCTCTATGGCGGAAGCGGGTATCAC-3’ and the reverse primer 5’-TCTAGAGTCGCGGCCGCTTATTTAACTTTGGTGAAGGTGTCCTGACCCT-3’. The TfRΔecto-RFP template was amplified using the forward primer, 5’-ACACCTTCACCAAAGTTAAATAAGCGGCCGCGACTCT-3’, and the reverse primer, 5’-ATACCCGCTTCCGCCATAGagGCGCCGGTGGAGTG-3’. Both fragments underwent Dpn1 digestions to remove template DNA. After template removal, both fragments were combined via Gibson assembly (New England Biolabs), and the resulting TfRΔecto-RFP-mSA clone was verified by sequencing.

Finally, the RFP was replaced by mEGFP using Gibson Assembly. The plasmid encoding mEGFP (alanine to lysine mutation at the 206^th^ amino acid to prevent dimerization) was generously provided by Dr. Adam Arkin (University of California-Berkeley). The mEGFP fragment was isolated with the forward primer 5’-GTAAAGGGGATCCACCGGTTATGGTGAGCAAGGGCG-3’ and reverse primer, 5’-TCCTCGCCCTTGCTCACCATAACCGGTGGATCCCC-3’. Using TfRΔecto-RFP-mSA as a template, the vector fragment lacking RFP was amplified using the forward primer 5’-GCATGGACGAGCTGTACAAGTCTATGGCGGAAGCGGGTATCAC-3’, and the reverse primer 5’-ATACCCGCTTCCGCCATAGACTTGTACAGCTCGTCCATGC-3’. Both fragments underwent Dpn1 digestions to remove template DNA. After template removal, both fragments were combined via Gibson assembly to generate TfRΔecto-mEGFP-mSA, as verified by sequencing.

### Cell culture and transfection

SUM159 cells were gene-edited to contain a HaloTag on both alleles of the AP-2 σ2 domain. These cells were generously provided by the Kirchhausen laboratory at Harvard University. These cells were grown in media composed of a 1 to 1 ratio of DMEM high glucose and Ham’s F-12 which were both purchased from Cytiva, pH 7.4. The media was supplemented with 5% fetal bovine serum (Cytiva), 10 mM HEPES (ThermoFisher Scientific), 5 μg-mL^-1^ insulin (Sigma-Aldrich), 1 μg-mL^-1^ hydrocortisone (Sigma-Aldrich), and 1% penicillin/streptomycin/L-glutamine (Cytiva). Cells were maintained at 37°C, 5% CO_2_. Cells were seeded onto acid-washed coverslips 24 hours before transfection at a density of 1×10^5^ cells per well in 6-well plates. Transfection was performed with 1 μg of plasmid DNA in combination with 3 μL of Fugene HD transfection reagent per well (Promega).

HaloTagged AP-2 was visualized in the SUM159 cells through the addition of membrane-permeable JaneliaFluor646-HaloTag ligand (Promega) at a concentration of 125 nM, 15 minutes prior to imaging. Lipid concentration before imaging was estimated using a nanodrop (ThermoFisher Scientific) to measure the absorption of Texas Red. Texas Red-DHPE was present at a 1 to 10 molar ratio within liposome mixtures. 1 μM of total lipid was incubated with the cells for 15 minutes at 37°C before imaging. The cells were washed with fresh phenol-red-free media containing more liposomes at a similar concentration and then imaged immediately.

### Preparation of liposomes

Lipids were dissolved in chloroform and stored at –80°C. Aliquots were brought to room temperature and combined at the ratios stated in the main text. Once mixed, the lipids were dried using a gentle stream of nitrogen. The remaining lipid film was dried for a minimum of 3 hours under vacuum. The lipid film was hydrated and thoroughly mixed into pH 7.4 buffer containing 25 mM HEPES and 150 mM NaCl. The lipid film was allowed to hydrate and swell on ice for 30 minutes. Liposomes were made via probe tip sonication using a Branson Ultrasonics SLPe Sonicator. The average liposomes diameter was measured using dynamic light scattering, and ranged from 70-80 +/-15 nm, as shown in Supplementary Figure 1.

### Total internal reflection fluorescence (TIRF) microscopy

TIRF microscopy was used to image live cells over the course of 10 to 12.5 minutes at 2.25-second intervals between frames. The TIRF system used an Olympus IX73 microscope body, an Olympus 60×1.45 NA Plan-Apo oil immersion objective, an external THORLABS TL2X-SAP Super Apochromatic objective, a Photometrics Evolve Delta EMCCD camera, and Micromanager version 2.0.0-γ1. The slide that mounted the coverslip was heated to 37°C using a microprocessor-controlled, home-built slide heating system. The TIRF system used 473 nm, 532 nm, and 640 nm lasers. Live-cell imaging occurred approximately 17 hours after transfection in phenol red-free media containing the equivalent of 1 μM of total lipid. Imaging media also contained OxyFluor (Oxyrase) at a ratio of 1 μL OxyFluor per 33 μL of imaging media.

### Tethering of liposomes

Liposomes were tethered using a method described previously^54^. No. 1.5H glass coverslips (Thor Labs) were cleaned using a 2% v/v Hellmanex III (Hellma Analytics) solution. Similarly, 4mm thick silicone gaskets were cleaned using 2% v/v Hellmanex III as well. The silicone gaskets contained 10 mm diameter holes and were washed thoroughly using ultrapure water and dried under a nitrogen stream. Placement of the gasket onto the cleaned coverslip created a tight seal to create an imaging well. The exposed coverslip within the imaging well was passivated using a layer of biotinylated polylysine-PEG-5kDa (PLL-PEG). PLL-PEG was created by combining a 49:1 molar ratio of PEG-to-PEG-Biotin. The PEG combination was mixed into a 20 mg/mL solution of PLL in 50 mM sodium tetraborate, pH 8.5. This mixture was continuously stirred at room temperature overnight and was then buffer exchanged into 25 mM HEPES, and 150 mM NaCl, pH 7.4, using 7 kDa molecular weight cutoff Zeba size exclusion columns (ThermoFisher). To passivate the glass, 10 μL of PLL-PEG was added to each empty gasket, allowed to incubate for 20 minutes, then serially rinsed using 25 mM HEPES, 150 mM NaCl, pH 7.4, and slowly pipetted into the well until at least a 15,000× dilution was achieved. Then, 2 μL of a NeutrAvidin solution consisting of 4 μg NeutrAvidin (ThermoFisher) dissolved in 25 mM HEPES, 150 mM NaCl (pH 7.4) was added to the passivated well and allowed to incubate for 20 minutes. Wells were similarly rinsed using the same buffer until a 15,000× dilution was achieved to remove unbound NeutrAvidin. Sonicated liposomes containing 0, 10, or 20 mol% 18:1 Biotinyl Cap PE (Avanti Polar Lipids) were added to the wells at a 1 μM total lipid concentration. Liposomes were incubated with the coverslip for 15 minutes, prior to washing with the same buffer until a 15,000× dilution was achieved, to remove excess liposomes.

### Calibration of liposome diameter

Liposomes were tethered to passivated coverslips as described above. Images of these liposomes were taken with a minimum of 15 acquisitions per well. These movies were analyzed using CMEAnalysis^51^ to acquire the maximum intensity over the local background for liposomes that are present in at least 3 simultaneous frames. From these data, the distribution of liposome fluorescence intensities was determined. The distribution of liposome diameters was measured using dynamic-light scattering and was converted to a distribution of liposome surface areas, assuming that all liposomes were approximately spherical. The distribution of intensities was compared to the distribution of surface areas to determine a conversion factor between surface are and intensity. Using this conversion factor, the intensity of liposomes beneath cells could be used to estimate the liposome surface area and diameter.

### Image analysis

Tethered liposomes and cell images were both analyzed using open-source detection software, CMEAnalysis, previously described by Aguet et al^51^. CMEAnalysis fits each fluorescent punctum with a two-dimensional Gaussian to the fluorescence intensity profile to each diffraction-limited punctum. Tethered liposomes had to be present in the first 3 frames of a short-time series to be considered valid. For cell movies, the center gaussian fits of the master channel were identified and subordinate channels were allowed to shift up to 3 standard deviations from the center of the master channel. Data were filtered according to the “significant-master” criterion assigned to each subordinate channel. This criterion is described by Aguet et al., but briefly states that a subordinate channel is positive for significant master, meaning it could be tracked itself if it colocalizes over a statistically significant number of frames, where the interaction through time is not due to chance. The internalization criterion was a custom-built MATLAB filter available upon request.

### Internalization Filter

To determine if a liposome was truly internalized, the liposome channel was tracked as the primary channel and checked for colocalizations with the AP2 channel. The primary channel is the channel that is tracked over time by the software and must be present throughout the track, whereas subordinate channels may colocalize for all or part of a track with the primary channel. Using a custom MATLAB script, we compared the signals of the primary (liposome) and subordinate (AP2) channels, applying a series of filters to identify true internalization events. The first filter criterion was that the liposome punctum had to be colocalized with an AP2 punctum across a statistically significant number of frames. This threshold was identified after determining the probability of random colocalization of the primary and subordinate channels. Using this threshold, we only retain tracks for which the colocalized duration is long enough to provide 95% or greater confidence of non-random colocalization. This type of threshold has been previously established in CMEanalysis^51^.

The second criterion that we applied identifies if a colocalized endocytic structure resides for a long enough period for the endocytic site to capture and internalize a liposome. To address that the endocytic structure was present for sufficient time, we required that at least 3 frames in the AP2 channel were at least 60% of the maximum tracked AP2 intensity. By using a frame rate of 2.25 seconds, we required that this intensity threshold is present for roughly 7 seconds or more in length which we empirically found to be true for observed internalized liposomes. By incorporating the intensity and temporal requirements we eliminate liposomes that might sample endocytic structures over short time scales but diffuse away prior to internalization.

The third criterion was designed to ensure that the liposome and AP2 signals disappeared from the TIRF field at approximately the same time. Specifically, the filter required the AP2 channel to display a drop of at least 50% of its maximum intensity within two frames of the end of the analyzed liposome track. By testing within 2 frames, we have allowed for a buffer of up to 5 seconds where the AP2 signal can depart and still count as a simultaneous departure. This is due to the intracellular nature of the AP2 signal which could disappear from the evanescent field prior to the liposome, and from the chromatic aberration of entering the TIRF field, which can lead to slightly different TIRF penetration depths between the two fluorescent channels.

The fourth criterion mandated that there must not be any additional drops in the AP2 signal other than the one present in the third criterion. The lack of additional drops had to be true from the time of the third criterion drop to the end of the liposome track, which contained 5 added buffer frames at its end. This requirement ensures that the liposome has truly departed, rather than simply dissociated from the endocytic site.

The last criterion states that any endocytic site that meets the previous requirements must only possess a signal lower than 25% of the maximum tracked AP2 intensity within the buffer frames. This criterion throws out AP2 signals that may be too close to the noise threshold, which would erroneously classify a liposome that meets criteria one through four as an internalized liposome, when in fact it is associated with a transient fluctuation in the AP2 signal but may have been internalized by a different pathway.

### Statistics and Plotting

Students T-tests, Z-tests for means, and ANOVA analysis were all run with α = 0.05, in either Microsoft Excel or Origin. Plotting was performed in the two aforementioned programs as well as MATLAB. Cartoons were made using the assistance of Adobe Illustrator and BioRender.

### Author Contributions

G.A.A. designed and performed experiments, analyzed the resulting data and wrote the manuscript. K.K., and S.G. performed experiments. J.R.H., and C.C.H. designed experiments and analyzed data. J.C.S. designed experiments and oversaw the research team. All authors consulted together on the interpretation of the results and the preparation of the manuscript.

## Acknowledgments

We thank Dr. Thomas Kirchhausen (Harvard University) for generously providing the SUM159 cell line containing the gene-edited AP-2 σ2-HaloTag. This research was supported by the National Institutes of Health Grant R35GM139531 to J.C.S., the NSF DMR through grant 1934411 (P.R. and J.C.S), the Welch Foundation through grant F-2047 (J.C.S), the NSF grant 2218467 (B.B. and J.C.S), and the continuing fellowship from The University of Texas at Austin Graduate School to G.A.A.

## Supporting Material

**Supplemental Figure 1.**
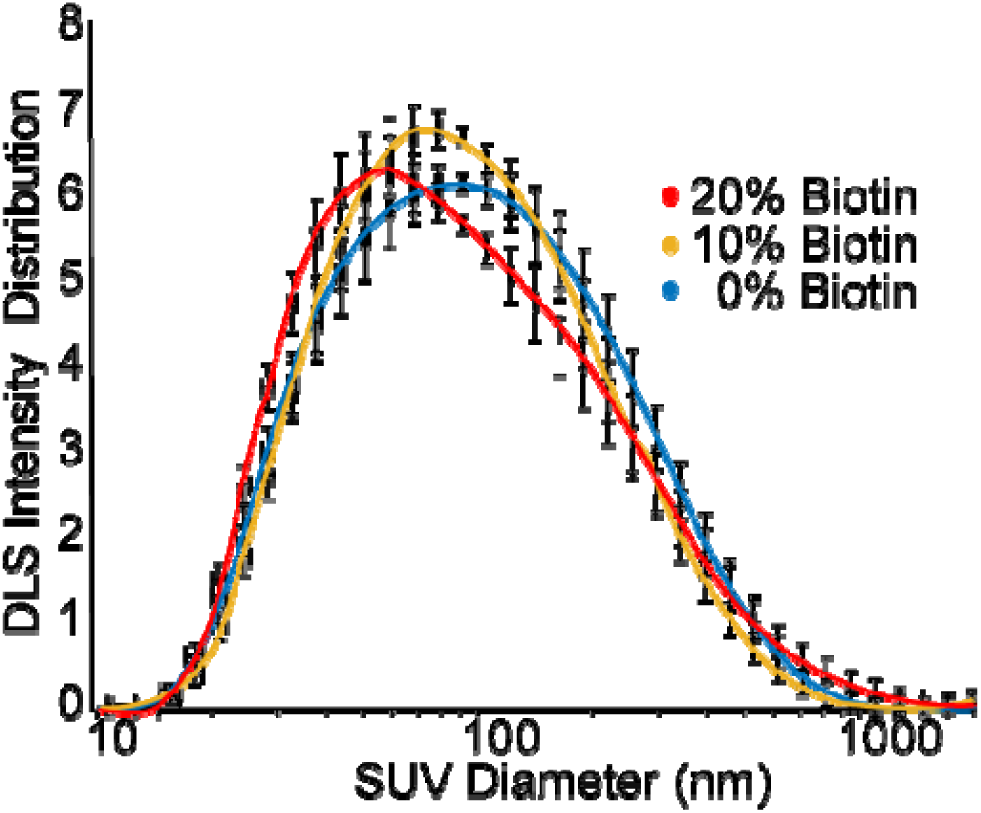
Diameter distributuions for liposomes containing either 0, 10, or 20 mol% biotinylated lipids. Each curve represents 6 independent trials measured by dynamic light scattering. The error bars represent the standard error of the mean for each group. ANOVA analysis of these distributions indicated that there were no significant differences between the groups.

**Supplementary Figure 2.**
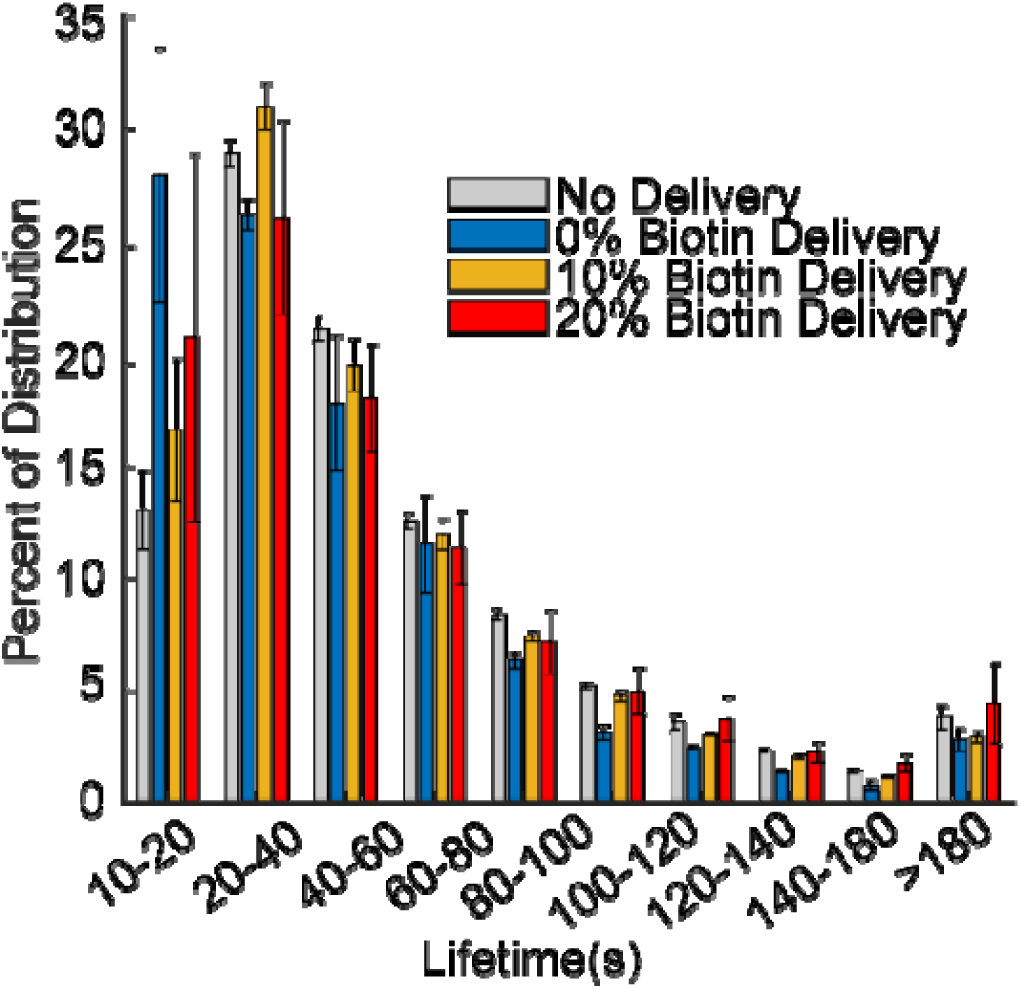
Distributuion of lifetimes for clathrin-coated structures in cell exposed to no liposomes (grey), and liposomes containing 0 (blue), 10 (gold), or 20 mol% biotinylated lipids (red). N = 3 independent trials for each trend. Bars represent the mean value within each lifetime cohort (horizatonal axis). Error bars are the standard error of the mean.

**Supplementary Video 1.** Example of a full cell containing JF646 labelled AP2 with SUM159 AP2-HaloTag cells interacting with Texas Red labeled liposomes containing 10% PE-CAP-Biotin. Liposome and endocytic machinery interactions were observed using TIRF microscopy.

**Supplementary Video 2.** Example of a single liposome interacting with a single clathrin-coated structure as it matures and is eventually internalized and cleaved from the membrane, disappearing into the cytosol.

